# Mutational switch-backs can accelerate evolution of *Francisella* to a combination of ciprofloxacin and doxycycline

**DOI:** 10.1101/2021.12.03.471061

**Authors:** Heer H. Mehta, David Ibarra, Christopher J. Marx, Craig R. Miller, Yousif Shamoo

## Abstract

Combination antimicrobial therapy has been considered a promising strategy to combat the evolution of antimicrobial resistance. *Francisella tularensis* is the causative agent of tularemia and in addition to being found in the nature, is recognized as a threat agent that requires vigilance. We investigated the evolutionary outcome of adapting the Live Vaccine Strain (LVS) of *Francisella* to two non-interacting drugs, ciprofloxacin and doxycycline, individually, sequentially, and in combination. Despite their individual efficacies and independence of mechanisms, evolution to the combination appeared to progress faster than evolution to the two drugs sequentially. We conducted a longitudinal mutational analysis of the populations evolving to the drug combination, genetically reconstructed the identified evolutionary pathway, and carried out biochemical validation. We discovered that, after the appearance of an initial weak generalist mutation (FupA/B), each successive mutation alternated between adaptation to one drug or the other. In combination, these mutations allowed the population to more efficiently ascend the fitness peak through a series of evolutionary switch-backs. Clonal interference, weak pleiotropy, and positive epistasis also contributed to combinatorial evolution. This finding suggests that, under some selection conditions, the use of non-interacting drug pairs as a treatment strategy may result in a more rapid ascent to multi-drug resistance and serves as a cautionary tale.

**Author summary:** The antimicrobial resistance crisis requires the use of novel treatment strategies to prevent or delay the emergence of resistance. Combinations of drugs offer one strategy to delay resistance, but the efficacy of such drug combinations depends on the evolutionary response of the organism. Using experimental evolution, we show that under some conditions, a potential drug combination does not delay the onset of resistance in bacteria responsible for causing tularemia, *Francisella.* In fact, they evolve resistance to the combination faster than when the two drugs are applied sequentially. This result is surprising and concerning: using this drug combination in a hospital setting could lead to simultaneous emergence of resistance to two antibiotics. Employing whole genome sequencing, we identified the molecular mechanism leading to evolution of resistance to the combination. The mechanism is similar to the switch-back route used by hikers while scaling steep mountains i.e., instead of simultaneously acquiring mutations conferring resistance to both drugs, the bacteria acquire mutations to each drug in alternating manner. Rather than scaling the steep mountain directly, the bacteria ascend the mountain by a series of evolutionary switch-backs to gain elevation and in doing so, they get to the top more efficiently.

## Introduction

The declining inventory of effective antibiotics is the direct result of a continuing rise of multi- drug resistant pathogens. It has been sensibly suggested that drug combinations that can inhibit multiple cellular targets rather than a single essential pathway can provide more successful antimicrobial therapies (1). This has accelerated the use of polypharmacological strategies in various therapeutic areas like antiviral therapies, treatment of nervous system disorders and cancer (2). In the field of antimicrobial resistance, this type of combination therapy has long been considered an effective resistance management strategy—although outcomes have produced very mixed results (3–6). Understanding which drug combinations will be effective, their interactions, their pharmacodynamics and the evolutionary strategies used by bacteria to acquire resistance are important parameters for the success or failure of antimicrobial therapies (7). In hospitals, however, the severity of multidrug resistant infections can drive doctors to empirically administer drug combinations, usually involving broad-spectrum agents (8). This strategy, although controversial, is recommended as a de-escalating strategy for severe infections (9, 10).

The success or failure of this empirical strategy critically depends on the physiological context, interactions between the drugs and the accessible evolutionary trajectories leading to resistance (7, 11). The likelihood that bacteria acquire resistance to a combination of drugs by adaptive mutations is dependent on several well-known parameters including mutation supply, population size, fitness effects and, perhaps most importantly, how the population effectively experiences antibiotic selection (1, 12). Administration of two drugs with distinct mechanisms of action would typically require at least two distinct adaptive mutations to arise simultaneously (7), which is significantly more difficult than a single adaptive change in response to a single drug. The simultaneous acquisition of multiple adaptive mutations within a single round of replication should be the product of their individual mutational frequencies (3), making it a rare event within a microbial population of a size relevant to a clinical setting (12, 13). This suggests that combination therapy with distinct classes of antibiotics should extend the efficacy of both drugs and greatly extend the time it takes to acquire resistance. However, many studies and clinical survey data have shown that, contrary to expectation, combination therapies can fail (4,5,14,15). For example, in the case of combination therapy for treatment of HIV infections, mismatched penetration of individual drugs into different body compartments can create zones of spatial monotherapy that can favor fast step-wise accumulation of resistance mutations, making the combination therapy ineffective (16). In the case of synergistic drugs, a single mutation can sometimes be sufficient to confer resistance to both drugs (3,17–19).

Thus, it is important to consider drug combinations that exploit evolutionary dynamics to maximize effectiveness of treatment rather than fail because of them (20). *Francisella tularensis,* the causative agent of tularemia, is an intracellular pathogen and an organism of concern because of its potential to be used as a bioterrorism agent (21). Medical management strategies in the case of exposure involve antimicrobial therapy, and stockpiles of ciprofloxacin (CIP) and doxycycline (DOX) are maintained by the CDC (Centers for Disease Control and Prevention) for treatment in mass casualty settings (22). Doxycycline is a second generation tetracycline that inhibits bacterial translation by reversible binding to the 30S ribosomal subunit (23, 24). Ciprofloxacin is a quinolone that inhibits bacterial replication by binding to DNA gyrases and topoisomerase IV (25–27). The importance of the use of these two drugs in a mass casualty setting underscores the need to understand the efficacy of these drugs and the evolvability of resistance to this combination. Both drugs have distinct mechanisms of action that target distinct cellular processes. Intuitively, we expect this combination to be effective because the evolution of resistance requires the acquisition of distinct mutations.

Contrary to expectation, we found that *in vitro* evolution of *Francisella tularensis* subsp. *holarctica* Live Vaccine Strain (LVS) to a combination of the non-interacting drug pair, CIP and DOX, occurred as quickly as evolution to one of the drugs individually. This, in effect, made the combination potentially worse than the individual drugs by simultaneously diminishing efficacy of both drugs. Surveys of the longitudinal order of mutations during adaptation of LVS to both drugs highlighted an interesting evolutionary mechanism. Rather than simultaneously acquiring mutations to both drugs, populations exposed to the drug combination developed resistance to each drug in an alternating but step-wise progression. We refer to this alternating evolutionary trajectory ascending the fitness landscape as “switch-backing” since it is reminiscent of a switch-back path ascending a mountainside. A straight path to the peak (analogous to simultaneous acquisition of mutations to both drugs), while being the shortest path, is not easily navigable. In this case, the opening move during evolution to the combination within all the replicate populations happened to be a mutation that conferred a marginal benefit to both drugs. The next mutation was an efflux pump mutation that helped with export of DOX, followed by a gyrase mutation that increased resistance to CIP and finally a mutation in the 30S ribosomal subunit that further increased resistance to DOX. The individual contribution of these mutations was resistance to one drug or the other, but their accumulation incrementally moved the evolving populations towards higher combinatorial resistance. The alternating pattern of mutation accumulation does for a population ascending a fitness peak what a switch-back route does for a hiker scaling a steep mountain.

If we are to achieve greater success in developing new combinatorial drug strategies, it is important to consider how drug combinations might exploit, or fall victim to, unexpected evolutionary dynamics. Our work illuminates how one such unexpected evolutionary strategy went on to defeat a potential combinatorial therapy and serves as a broader cautionary tale.

## Results

### The time for evolution of resistance to the two-drug combination was shorter than that of the individual drugs administered sequentially

*In vitro* experimental evolution was used to determine the evolvability of resistance to CIP and DOX. A checkerboard assay revealed that the two drugs, when used together on LVS, formed a non-interacting pair (Fractional Inhibitory Concentration Index (FICI) = 2) (FICI interpretation based on (28)), which is consistent with their known independent targets. Thus, we predicted that this combination could potentially extend the timeline for evolution of resistance since it would require the simultaneous acquisition of mutations to both drugs. Four different selection environments were investigated for the two drugs, CIP and DOX (Fig. 1). Five replicate populations were evolved in each of the selection environments.

i. Mono-selection: Ancestor LVS was evolved to (a) DOX or (b) CIP using a sub-inhibitory concentration gradient increasing in a step-wise manner. Replicate populations of LVS took 48 days to evolve resistance and grow at 8 µg/ml DOX and 33-35 days to grow at 1 µg/ml CIP (Fig. 2 A). These concentrations of DOX and CIP are two-fold higher than their clinical MIC breakpoint (29), indicating that the populations had acquired complete resistance at these drug concentrations.
ii. Sequential-selection: Mono-evolved populations were subsequently evolved to the second drug in the absence of the first selective agent. (a) DOX resistant populations evolved by mono-selection were further evolved to CIP and (b) CIP resistant populations were further evolved to DOX. The former took 32 days while the latter took an additional 28 days (Fig. 2 A). Despite the removal of the first selective agent, sequentially evolved populations and end point isolates obtained from these treatments continued to have resistance to both drugs (Fig. 2 B)
iii. Sequential-selection while maintaining a high concentration of drug 1: The highest dose of selective agent used in mono-selection was maintained while mono-evolved populations were subsequently evolved to the second drug: (a) DOX resistant populations evolved by mono- selection were subsequently evolved to CIP in the presence of 8 µg/ml DOX throughout sequential selection ; (b) CIP resistant populations were evolved to DOX in the presence of 1 µg/ml CIP throughout sequential evolution. The former took 37 days while the latter took an additional 30 days (Fig. 2 A).
iv. Combination selection: Ancestor LVS was evolved to a combination of DOX and CIP applied in a ratio that was proportional to the ratio of their individual MICs using a step-wise gradient of sub-inhibitory concentrations. Replicate populations evolved to the drug combination in 51-54 days (Fig. 2 A).

**Figure 1:**
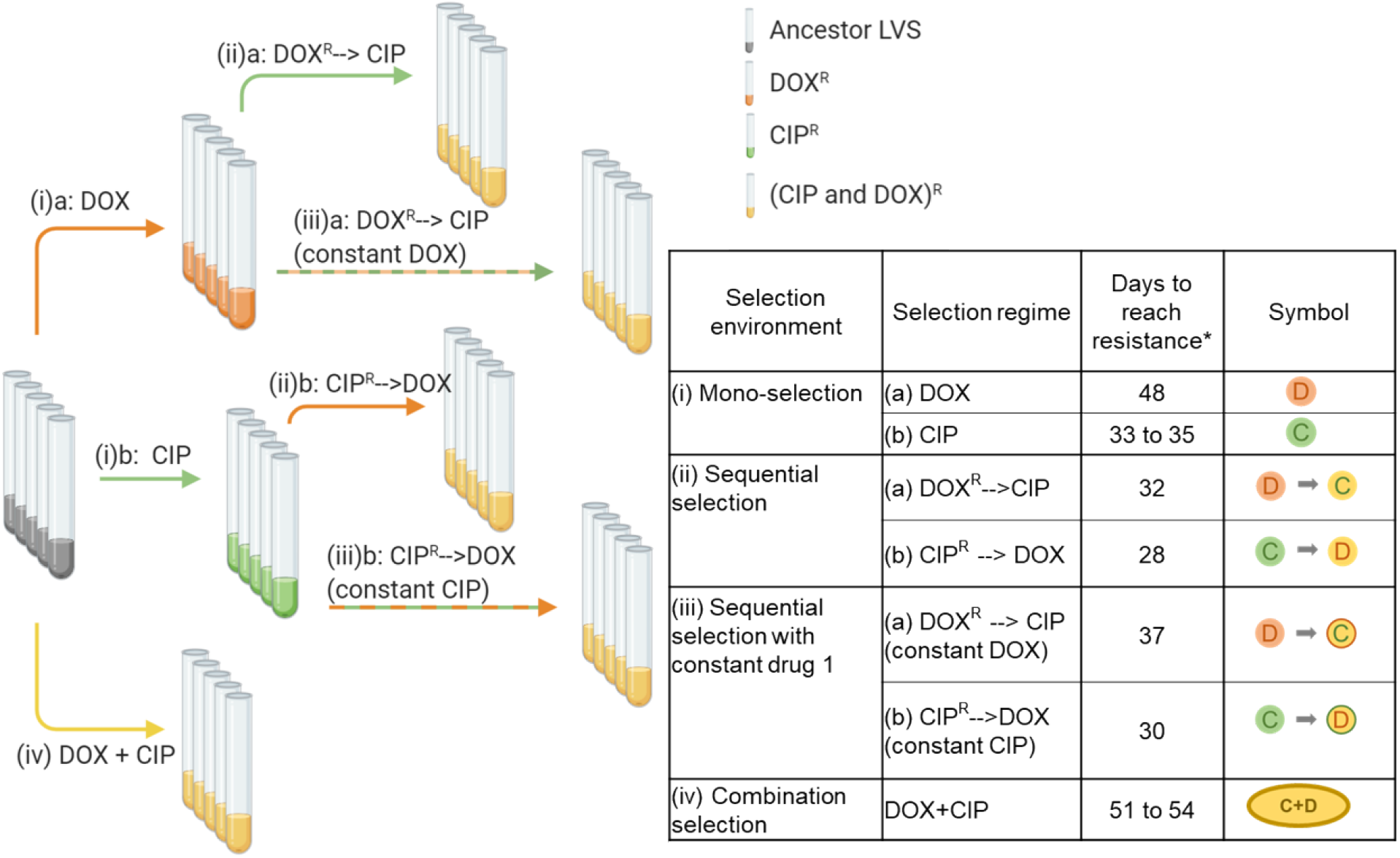
*In vitro* experimental evolution of LVS to CIP and DOX. The figure on the left shows the scheme of evolution with the various selection regimes and the table on the right shows the four selection environments used, the different regimes for DOX and CIP within each environment, the number of days taken to evolve populations within each regime as well as the symbols used to identify each condition. *Five replicate populations were set up for each selection regime. Created with BioRender.com.

**Figure 2:**
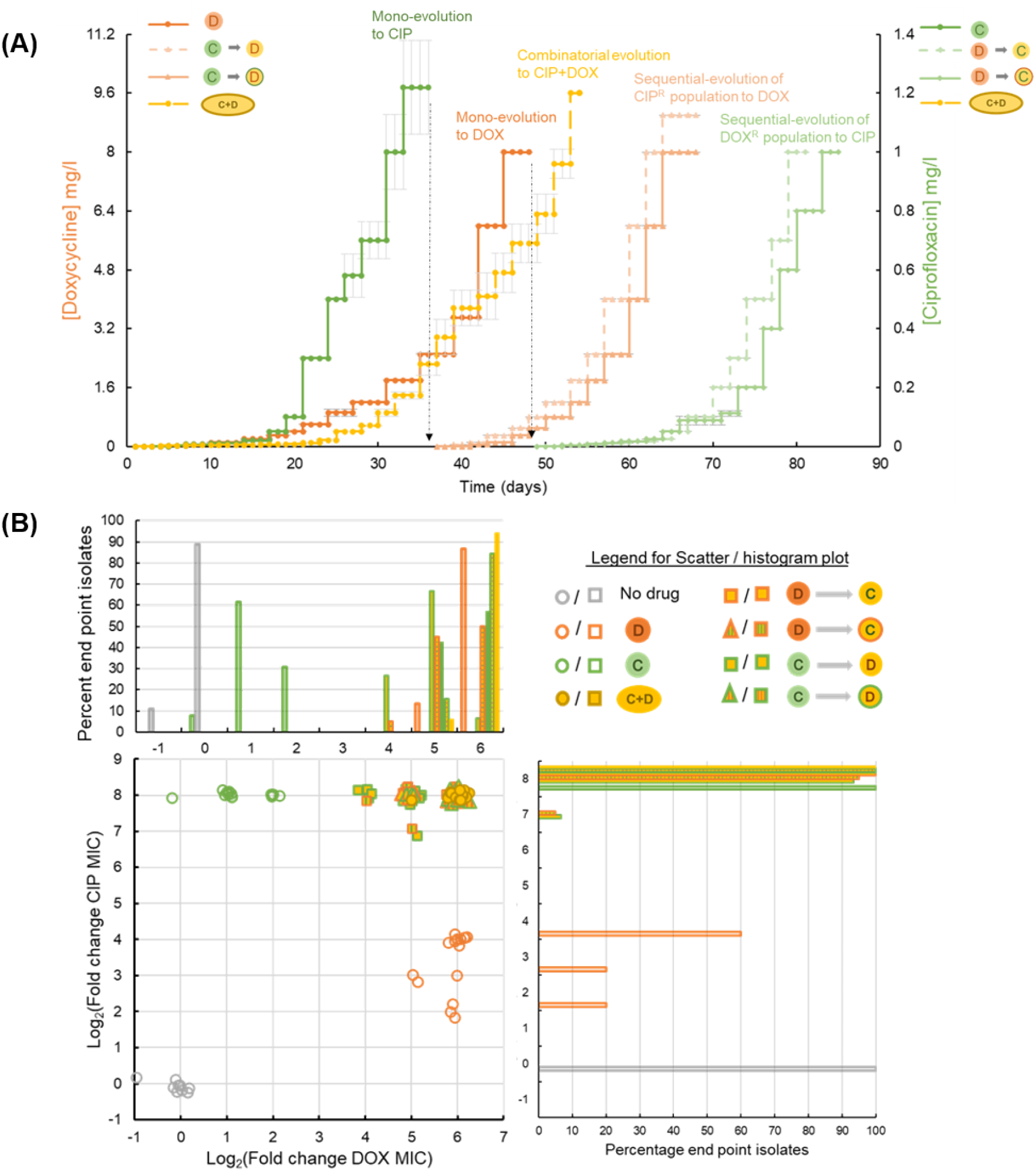
(A) Timeline of evolution to different selection environments. The graph shows the step-wise gradient of sub-inhibitory drug concentrations used during evolution of LVS. Text at the top of each curve indicates the selection condition used. The primary Y-axis (orange) represents the DOX concentration in mg/l and the secondary Y-axis (green) represents the CIP concentration in mg/l. Symbols for the various selection conditions used are from Fig. 1. The same symbols are shown alongside each of the Y-axes to indicate the axis (or the drug) relevant to each curve. For the yellow curve, which represents timeline of evolution to combinatorial selection, DOX concentrations can be seen on the left and CIP concentrations on the right. The black dotted line indicates the use of the relevant mono-evolved population for sequential selection to the second drug. (B) Distribution of evolved end point isolates based on their Minimum Inhibitory Concentrations (MICs) of DOX and CIP. The X and Y axes on the scatter plot represent the log_2_ fold change in MIC of DOX and CIP compared to ancestor. Noise has been added to each data point to create separation. The mono-evolved end point isolates cluster in independent regions indicating resistance to the selection used, while all the sequential and combinatorial evolved end point isolates cluster in the top right corner of the graph which indicates high levels of resistance to both drugs. The histogram above the scatter plot shares its X-axis with the scatter plot and the one on the right shares its Y-axis with the scatter plot. Each histogram shows the percentage of end point isolates from each selection regime falling into the respective log_2_ MIC fold change category.

Appropriate controls were run for each selection regime which involved evolving the ancestor LVS to non-selective media, evolving the mono-evolved populations to non-selective media and evolving the mono-evolved populations to a constant dose of the first selective agent.

The timeline to resistance via mono-selection and sequential selection matched our expectation that the two drugs worked via independent mechanisms and thus, mutations conferring resistance to mono-evolved populations were unable to potentiate complete resistance to the second drug, causing sequential selection to take an additional 28 to 37 days. However, phenotypes of the mono-evolved isolates did indicate some low-level pleiotropy as DOX evolved isolates had acquired a slightly improved ability to tolerate CIP and vice-versa (Fig. 2 B and Table S1). Removal of the selective pressure from the first selective agent did not lead to loss of resistance, suggesting that fitness cost associated with resistance to either drug was low. Fig. 2 B shows the MICs of CIP and DOX for end point isolates obtained from the different selection regimes.

**Table 1:** Strains and plasmids

| Strain or plasmid | Description | Reference or source |
| --- | --- | --- |
| Strains |  |  |
| LVS | *Francisella tularensis* subsp. *holarctica* live vaccine strain | Dr. Karen Elkins |
| FupA/B R343I | LVS *fupA/B*<sup>1028G→T</sup> | This work |
| TolC I436T | LVS *tolC*<sup>1307T→C</sup> | This work |
| GyrA D87Y | LVS *gyrA*<sup>259G→T</sup> | This work |
| S10 V59L | LVS *FTL_0235*<sup>175G→T</sup> | This work |
| LVS-2X | LVS *fupA/B*<sup>1028G→T</sup> *tolC*<sup>1307T→C</sup> | This work |
| LVS-3X | LVS *fupA/B*<sup>1028G→T</sup> *tolC*<sup>1307T→C</sup> *gyrA*<sup>259G→T</sup> | This work |
| LVS-4X | LVS *fupA/B*<sup>1028G→T</sup> *tolC*<sup>1307T→C</sup> *gyrA*<sup>259G→T</sup> *FTL_0235*<sup>175G→T</sup> | This work |
| Plasmids |  |  |
| pMP812 | Km<sup>R</sup> suicide vector | (45) |
| pMP-fupR343I | Km<sup>R</sup> pMP812 suicide vector bearing a 2kb fragment containing nt. position 1028 (SNP G → T) of gene *fupA/B* | This work |
| pMP-tolCI436T | Km<sup>R</sup> pMP812 suicide vector bearing a 2kb fragment containing nt. position 1307 (SNP T → C) of gene *tolC* | This work |
| pMP-gyrAD87Y | Km<sup>R</sup> pMP812 suicide vector bearing a 2kb fragment containing nt. position 259 (SNP G → T) of gene *gyrA* | This work |
| pMP-S10 V59L | Km<sup>R</sup> pMP812 suicide vector bearing a 2kb fragment containing nt. position 175 (SNP G → T) of gene *FTL_0235* encoding 30S ribosomal protein S10 | This work |

Replicate populations of the strain were able to evolve total resistance to the drug combination in 51-54 days, which was only 6 days more than it needed to acquire resistance to DOX alone. As LVS grows slowly, this corresponded to just 12-18 additional generations. Compared to sequential evolution, which took a total of 65 to 85 days, combinatorial evolution did not extend the time taken to develop resistance to the drug combination. The rapid adaptation to the drug combination was surprising since synergy was not observed in the ancestor and it was anticipated that any adaptive changes resulting in resistance would be, of necessity, independent of each other, owing to their well-established and independent mechanisms-of-action and known cellular targets as well as timeline for sequential selection.

### Sequentially adapted populations acquired the same class of mutations as mono-evolved populations

Whole genome sequencing of resistant populations and randomly selected end-point isolates obtained from replicate populations identified mutations that had been acquired during adaptation. These mutations could be classified into 4 major systems: (i) transporter FupA/B (involved in iron transport (30)) (ii) DNA gyrase/topoisomerase IV (targeted by CIP (27)) (iii) tripartite efflux pump AcrAB-TolC (involved in efflux of antimicrobials (31)) (iv) 30S ribosomal subunits (targeted by DOX (24)). Figure 3 A shows a heat map of the types of mutations identified among end point isolates from each selection condition.

**Figure 3:**
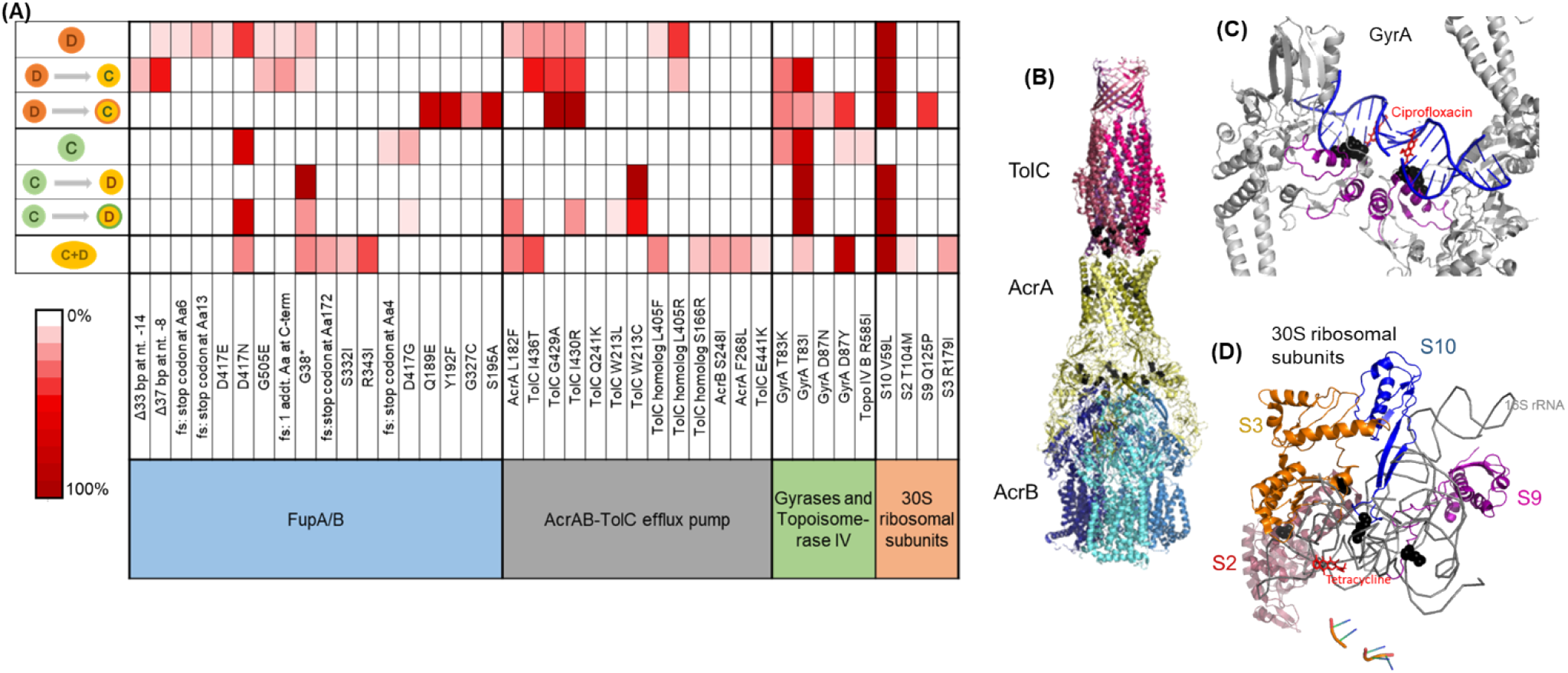
Mutation categories involved in resistance and distribution of mutant alleles among end point isolates obtained from different selection conditions. (A) Heat map shows the frequency of non-synonymous mutant alleles within a mutation category identified in end point isolates obtained from each selection condition. 2 to 4 end point isolates from each of the five replicate populations and seven selection environments were sequenced. Each row represents one condition and symbols for selection conditions are those used in Fig. 1. fs = frameshift, TolC homolog = FTL_1107 (B) Mutations observed in AcrAB-TolC in adapted LVS isolates mapped onto the protein structure for this efflux pump from *E. coli* (PDB: 5V5S). Positions of mutations are indicated (black spheres). Most mutations are proximal to subunit interfacial contact regions that are implicated in allosteric control of the effluxer (35). (C) Mutations observed in GyrA in adapted LVS isolates mapped onto the protein structure of GyrA from *S. aureus* bound to ciprofloxacin (PDB: 2XCT). Adaptive mutations are directly proximal to the ciprofloxacin binding pocket of the gyrase (black spheres). (D) 30S ribosomal subunits S2, S3, S9 and S10 from *T. thermophilus* interacting with tetracycline (PDB: 1HNW). Mutant residues identified in LVS adapted strains are mapped onto this structure and are shown in black spheres. They are mostly found in loop regions interacting with the 16s rRNA that directly comprises part of the tetracycline binding pocket of the 30S ribosome.

As shown in Fig. 3 A, mono-evolution to CIP was a result of mutations in FupA/B and gyrases whereas mono-evolution to DOX was driven by mutations in FupA/B, AcrAB-TolC and 30S ribosomal proteins. While it was not surprising to observe mutations in the gyrases, 30S ribosomal proteins and efflux pumps in response to DOX and CIP, mutations in the iron transporter, FupA/B (30)—something that occurred in both mono-selection conditions—was unanticipated. Mutations in DNA gyrases (Fig. 3 C) were consistent with previous observations of mutations ocurring in the quinolone resistance determining region (QRDR) of the enzyme that offered protection from quinolones like CIP (32). Mutations in the 30S ribosomal subunits (Fig. 3 D), specifically subunit S10 have been linked to resistance to tetracyclines, of which DOX is a member (33, 34). AcrAB-TolC is a tripartite multidrug efflux pump. Deletion of *tolC* or its homolog, *FTL_1107* in LVS leads to increased susceptibility to many antibiotics (31). Mutations in this efflux pump specifically clustered close to the intermolecular subunit interfaces (Fig. 3 B) that are implicated in conformational changes during the transition from resting to the transport state (35). The pump is a highly allosteric system in which re-packing of protein-protein interfaces plays a role in opening of the TolC channel (35). We speculate that the mutations seen in adapted LVS isolates may allow increased recognition and subsequent efflux of DOX, since these mutations are observed only in DOX evolved isolates. FupA/B has been studied as a high-affinity transporter of ferrous iron and siderophore mediated ferric iron in LVS (30, 36). A structure of FupA/B was not available for mapping the adaptive mutations, but five out of the 18 mutant alleles identified in this gene (including the introdution of a stop codon at Gly-38) would be predicted to produce an early truncation of the protein (Fig. 3 A), consistent with a loss of function. No mutations in these four classes were observed in populations evolved in the no-drug control. Furthermore, mutations that commonly appeared in the no-drug control populations as well as the populations undergoing drug selection were removed from further investigation since they were most likely mutations associated with adaptation to the growth media. Concomitantly, none of the isolates from the no- drug control evolved populations had acquired any increase in resistance to CIP or DOX (Fig. 2 B).

During sequential selection, we observed new mutations without the loss of previous mutations acquired during mono-selection, thereby making sequentially evolved populations resistant to both drugs, whether or not a constant dose of the initial drug was present during sequential selection. Evolution of CIP resistant populations to DOX was accompanied by mutations in the AcrAB-TolC efflux pump and 30S ribosomal subunits while DOX resistant populations became CIP resistant by acquiring mutations in the gyrases (Fig. 3 A). These findings were consistent with the initial premise that resistance to the two drugs is mediated through distinct evolutionary trajectories. Although the two drugs were not synergistic in this organism, evolution to each of the drugs did include mutations in FupA/B, which may explain why isolates evolved by mono-selection to DOX also acquired low levels of cross-resistance to CIP and vice versa (Fig. 2 B). Mutations implicated in resistance identified in end point isolates are shown in tables S1-S4. Datasets S1-S5 contain the complete list of mutations in end point isolates and populations.

### Mutations in FupA/B decreased drug concentrations inside the cell but provided only a modest increase in resistance

Since the drugs used in this study target cytoplasmic machinery inside the cell, we reasoned that perhaps the mutations in FupA/B influenced antibiotic transport and not just its canonical role as an outer membrane iron transporter. To test this hypothesis, we conducted antibiotic accumulation assays and MIC assays on the ancestor and an engineered point mutant, FupA/B^R343I^, one of the alleles identified during *in vitro* evolution. Accumulation assays showed that the FupA/B mutant accumulated approximately 10-fold less tetracycline and 3-fold less CIP than the ancestor (Fig. 4 C), suggesting that this mutation reduced the rate of drug accumulation inside the cells. This was also consistent with the appearance of SNPs and frameshifts in *fupA/B* that caused early termination suggesting that inactivation of *fupA/B* decreases drug accumulation (Fig. 3 A). It should be noted that, for this assay, tetracycline is used as a proxy for DOX since tetracycline forms a yellow compound upon reaction with HCl and can be easily quantified by spectrophotometry (See methods). The MIC of tetracycline in the ancestor LVS strain and the FupA/B^R343I^ mutant is the same as that of DOX and thus, it makes a good proxy for DOX.

**Figure 4:**
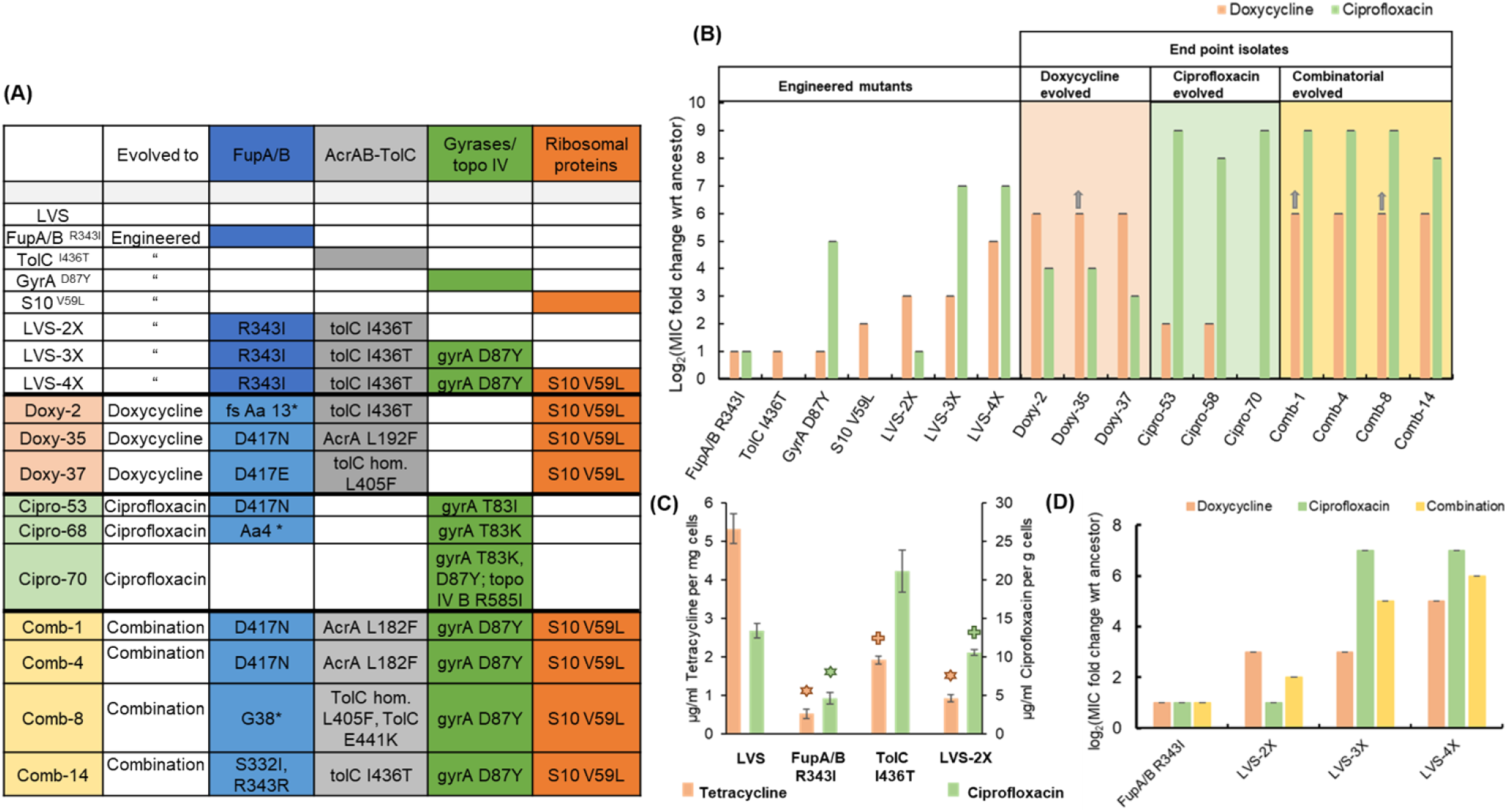
(A) Genotypes of engineered mutants and selected end point isolates obtained from evolution studies. The individual adaptive steps of the most successful evolutionary trajectory of replicate population 3 (shown in Fig. 5 A) was reconstructed via allelic exchange in the ancestor LVS strain. For end point isolates, mutant alleles are only shown for genes implicated in resistance (full genotypes are included in DataSets S1-S5). (B, D) MIC values for the strains representing reconstructed steps toward adaptation (engineered mutants) and selected end point isolates are shown as log_2_ fold change with respect to MIC of ancestor strain. Bars indicate average of 3 biological replicates with error bars showing standard deviation. Grey arrows indicate that MIC value is greater than the highest antibiotic concentration tested. (C) Tetracycline and ciprofloxacin accumulation assay: Tetracycline (used as a proxy for doxycycline) accumulated in the wild type and mutant cells is denoted by orange bars on the primary Y-axis and ciprofloxacin accumulated is denoted by green bars on the secondary Y-axis. Accumulated drug concentrations have been normalized by the wet weight of the cell pellets. Bars indicate average of 3 biological replicates and error bars denote standard deviation. Symbols above the bars denote statistical significance compared to ancestor LVS (stars = p value < 0.005 and cross = p value < 0.05; two-tailed p value determined using Student’s t test for two samples assuming unequal variance).

The FupA/B^R343I^ mutant also had a modest, 2-fold increase in MIC of both, DOX and CIP (Fig. 4 B). While mutations in FupA/B can be classified as “generalist” mutations that offer protection from both drugs in this study, the degree of protection offered by this individual mutation is very modest.

Since the pump AcrAB-TolC is involved in efflux of antibiotics, the combined effect of mutations in FupA/B and AcrAB-TolC, seen in the DOX evolved isolates, on drug accumulation was also tested. While the single mutant TolC^I436T^ accumulated 2.5-fold less tetracycline than the ancestor, it showed no decrease in CIP accumulation, consistent with the observed increase in the MIC of DOX alone (Fig. 4 B). The double mutant encoding FupA/B^R343I^ and TolC^I436T^ (LVS 2X) was able to produce an 8-fold increase in MIC of DOX and consequently, a strong effect on tetracycline accumulation but a weaker effect in the CIP accumulation assay consistent with the MIC of each drug (Fig. 4 B and C).

### Combinatorial evolution proceeded in a highly conserved and reproducible manner

The successful evolutionary trajectories of both the mono- and sequential selection studies were consistent with known biological functions of the mutational targets but did not provide explanation for the rapid adaptation that was observed in the combinatorial selection studies (Figs. 1 and 2). Although the pleiotropic FupA/B mutation could have been a one-step solution to the problem of simultaneous evolution to both drugs, it imparted a very marginal (two-fold) improvement in resistance to the drug pair, which could not account for the 100-fold increase in resistance gained by the evolved population. In order to understand the underlying evolutionary dynamics that facilitated acquisition of resistance faster than anticipated, we examined the temporal population dynamics at a metagenomic level to quantitate the rise and fall of adaptive alleles over time (Fig. 5 A and B). This allowed us to trace the evolutionary trajectories that orchestrated the process of adaptation. All the mutational categories observed in the mono- and sequential-evolved populations were identified in populations undergoing combinatorial evolution. Furthermore, the order in which these categories arose in each population was highly conserved. Among the five replicate populations, there was consistent parallel evolution (Fig. 5 and Fig. S1). FupA/B mutations were the first ones to arise, typically when the drug concentration in the population was just below the MIC of the ancestor strain. Subsequently, mutations in AcrAB-TolC arose. The specialist mutations in the gyrases and the 30S ribosomal subunits appeared later once the population was growing at a drug concentration well above the ancestral MIC (Fig. 5 A and B).

**Figure 5:**
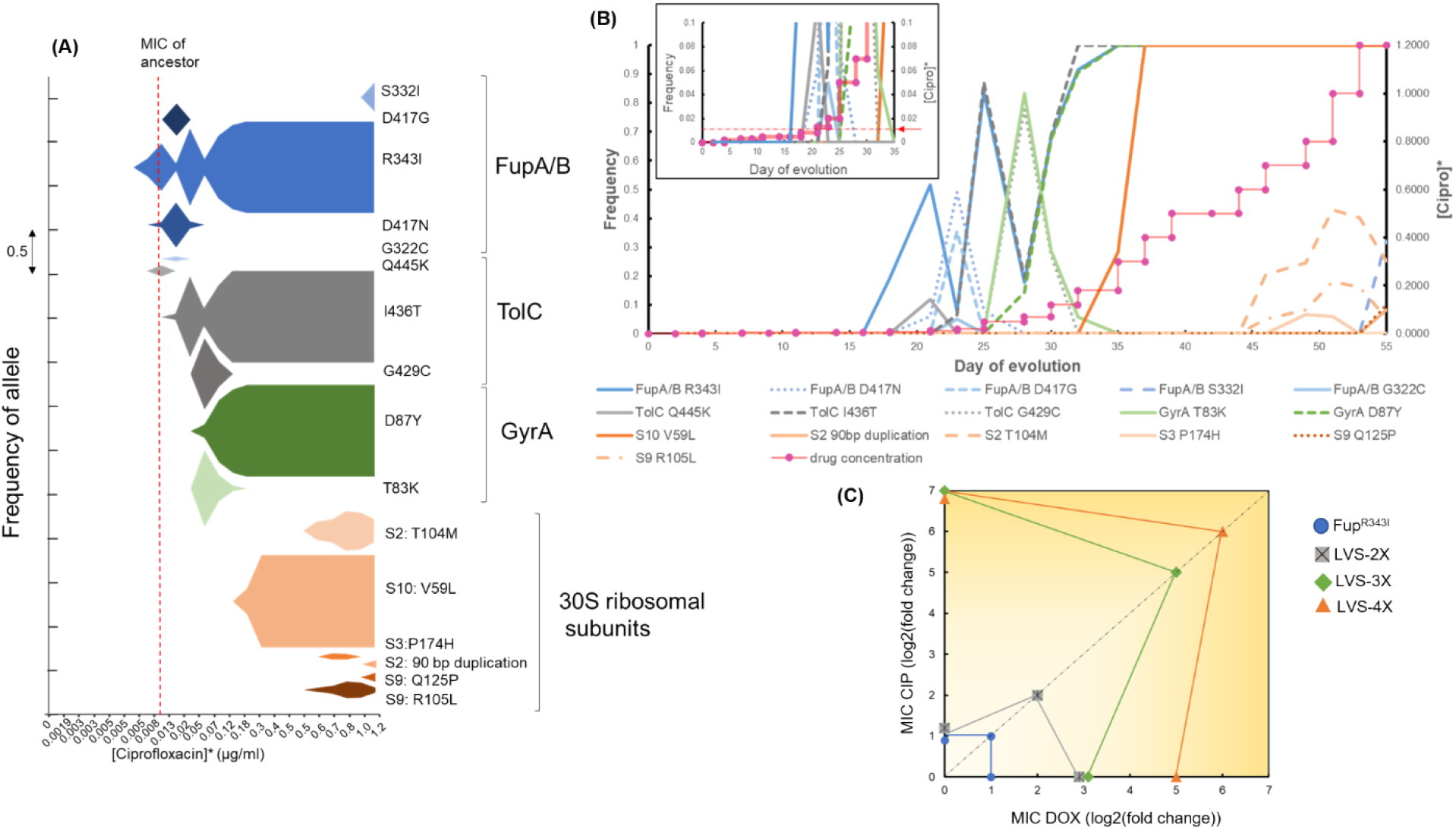
Order of mutations acquired during combinatorial evolution to CIP and DOX. (A) Metagenomic analysis of a single population as it adapted to both doxycycline and ciprofloxacin simultaneously. The x-axis is the drug concentration used during adaptation while the area of the plots is the frequency of the specific allele. *Ciprofloxacin and doxycycline were applied simultaneously such that doxycycline concentration was 8 times that of ciprofloxacin. The color shading corresponds to the general mutational class e.g. blue=FupA/B, gray=TolC, green= GyrA and orange=30S ribosomal proteins. It is clear from this figure that 4 mutant alleles, FupA/B^R343I^, TolC^I436T^, GyrA^D87Y^ and S10^V59L^ were the most successful in this population and reached saturation. These were the alleles selected to re-construct the mutations in the ancestor strain in the order in which they arose in this population. (B) Line graph showing the frequency of mutant alleles (on primary Y-axis) arising in the same population shown in A plotted against the day of adaptation. The secondary Y-axis shows the step wise gradient of the drug concentration used during combinatorial selection (*Doxycycline concentration used = 8 times [Ciprofloxacin]). The inset shows a magnified version of the first 35 days of adaptation and the dotted red line with the red arrow shows the MIC of the ancestor strain. (C) Mutational switch-backs shown as the accumulation of mutations during evolution. Resistance phenotypes of the engineered mutants that were created in the order in which the mutations appeared in the evolving population are shown. The X-axis represents the change in MIC of DOX, the Y-axis represents the change in MIC of CIP and the diagonal dotted line represents the change in MIC for the combination of CIP and DOX. Change in MIC is with reference to ancestor and is represented as log_2_(fold change). Noise has been added to overlapping data points on the X- and Y-axes to show separation.

Longitudinal mutational data revealed strong and recurrent clonal interference dynamics within the emerging adaptive alleles for FupA/B and GyrA. From Fig. 5 A, we observed that multiple mutant alleles in FupA/B evolved and competed in the population before fixation of FupA/B^R343I^. This was also observed in the early TolC and GyrA alleles. It was also interesting to observe the dynamics between the emerging FupA/B and GyrA mutations. Since FupA/B was consistently the first mutation to arise in all the replicate populations, it potentially played an important role in the process (Fig. S1). However, in three of the five evolving populations, FupA/B mutations dipped to very low levels around the same time the first gyrase mutations arose in the population (Replicate 2, 3 and 4 in Fig. S1). This drop in frequency was short-lived and FupA/B mutations managed to rise again and dominate the population. This type of clonal interference suggested that, at least temporarily, gyrase mutations were able to offer what Fup mutations had offered to the population. This was consistent with the MIC values for the engineered FupA/B^R343I^ and GyrA^D87Y^ mutants (Fig. 4 B). It is unclear why FupA/B mutations were able to rise again in these populations but it is clear that in spite of the clonal interference, ultimately, all four categories of mutations (shown in four different colors in Fig. S1) were needed for the populations to grow at the highest CIP and DOX concentrations used.

As an indifferent combinatorial pair, one might expect the accumulation of adaptive mutations to be equally indifferent i.e. the order of mutations and the systems they altered could have been random but this was clearly not the case. This reproducibility suggested that this order was likely an efficient path for the population to maneuver through the adaptive landscape and convert the challenge of simultaneously acquiring multiple mutations into a step-wise task, thus accelerating evolution of resistance.

### An evolutionary strategy of mutational switch-backs was validated by step-by-step re-creation of an evolutionary trajectory leading to combinatorial resistance

To investigate the significance of the order of appearance of mutations, allelic replacement was used to introduce successful mutant alleles from one of the replicate populations evolved to the combination of CIP and DOX (shown in Figs. 5 A, 4 A and 4 B) into the ancestor LVS strain. Mutations were introduced in the same order in which they arose during adaptation (See engineered mutants in Fig 4 A). The first step during adaptation was always a mutation in FupA/B, which, as discussed previously, served as a modest generalist. The FupA/B^R343I^ mutation in the LVS background conferred a two-fold increase in the MIC of CIP and DOX. Introduction of the next mutant allele, TolC^I436T^ into the FupA/B^R343I^ background produced the double mutant LVS- 2X which had an additional four-fold increase in DOX MIC but no change in susceptibility to CIP. Consistently, LVS-2X was more effective at reducing tetracycline than CIP accumulation (Fig. 4 B and C). This clearly suggested that mutations in FupA/B, while serving as a “generalist” during the evolution process, when combined with adaptive mutations within AcrAB-TolC, preferentially shifted the environment to weaken DOX selection pressure. Cells now experienced a relief from the stress imposed by DOX and the selection condition switched to mimic a mono-selection environment against CIP rather than the combination. In other words, the efflux pump mutations served as a switch-back for LVS during evolution.

At this stage, evolution progressed as if cells were being exposed to an alternating treatment of the two drugs rather than simultaneously. The next move made by the five replicate populations was to acquire mutations in the gyrases/topoisomerase IV genes, which are known to be specific targets of CIP. Introducing the GyrA^D87Y^ allele into LVS-2X (to make LVS-3X) dramatically raised the MIC of CIP to 64-fold above LVS-2X, while the DOX MIC remained at the same level as LVS- 2X (Fig. 4 B). This implies that the selection pressure shifted once again and DOX became the selective agent, representing another switch-back point. Finally, mutations in the subunits of the 30S ribosome arose, which raised the DOX MIC above the resistance breakpoint for this organism (29). The 30S ribosomal subunit mutation, S10^V59L^ was introduced into the LVS-3X strain to produce a 4X mutant that had a 32-fold higher level of DOX resistance compared to the ancestor.

Re-creating the genotype of the evolved mutant in a step-by-step manner allowed us to trace the evolutionary history of the cells and determine the contribution of each mutation during adaptation. An important observation from the engineered mutants was the unexpected phenotype of some strains. The large increase in resistance to CIP in the single GyrA^D87Y^ mutant was unsurprising, but it also conferred a two-fold increase in resistance to DOX (Fig 4 B). Introduction of GyrA^D87Y^ in LVS-2X (to make LVS-3X) did not improve resistance to DOX (Fig. 4 B) but conferred an eight-fold improvement in resistance to the combination of CIP and DOX (Fig. 4 D). Single mutants TolC^I436T^ and S10^V59L^ only improved resistance to DOX, not CIP when tested individually (Fig. 4 B). However, when these mutations were introduced in the background of FupA/B^R343I^ and LVS-3X, respectively, to create LVS-2X and LVS-4X, in addition to improving resistance to

DOX, they also led to a two-fold increase in resistance to the combination of CIP and DOX (Fig. 4 D). Thus, while individually these mutations protected against one of the two drugs (except GyrA^D87Y^), in combination with other mutations and in the presence of both, CIP and DOX, they appeared to show low levels of positive epistasis. As illustrated in Figure 5 C, over the course of evolution, switch-backs and low-level epistasis allowed the cells to quickly ascend a fitness peak leading to co-resistance.

## Discussion

The aim of this work was to explore the evolvability of LVS to a combination of CIP and DOX with the expectation that two drugs acting on distinct cellular processes would be effective in delaying resistance. To our surprise, LVS adapted rapidly to both drugs and highlighted an efficient evolutionary mechanism by which populations could acquire resistance through switch- backing. Unlike in other organisms (14, 37), in LVS, these drugs do not interact with one another and thus, the presence of one does not impact the ability of the other to inhibit growth. As such, resistance to this drug combination would be expected to require the simultaneous acquisition of mutations in both the targeted pathways. Not only did the combination not significantly delay the onset of resistance, but resistance was gained to both drugs on about the same timescale as either drug individually. Thus, a combinatorial therapy of DOX and CIP could actually make outcomes worse by simultaneously and quickly rendering both drugs impotent in some scenarios.

Mono-selection experiments demonstrated that, with the exception of mildly pleiotropic FupA/B mutations, resistance to both drugs was acquired via independent mechanisms, as expected. This was further supported by sequential selection experiments which demonstrated that, to become completely resistant to the second drug, additional mutations were necessary. It was apparent from the longitudinal study of adaptation to the drug combination that mutational switch-backs facilitated adaptation (Fig. 5). FupA/B mutations, as generalists, imparted low levels of resistance to both antibiotics and were the first to arise in the population. The specialist mutations known to be involved in resistance specifically to CIP (gyrases) and DOX (30S ribosomal S10) did not appear until the population was growing past the ancestral MIC (Fig. 5 A and B). The intermediate mutations in AcrAB-TolC provided increases in resistance to DOX and signified the first switch- back event. Figure 4 D shows that after the acquisition of the generalist FupA/B mutations, the sequence of events leading to the creation of LVS-4X involved a series of step wise changes that conferred an individual benefit to one drug or the other (LVS-2X showed improved DOX resistance, LVS-3X showed improved CIP resistance and LVS-4X showed further improved DOX resistance), but led to a gradual overall increase in resistance to the drug combination. Although FupA/B mutations were the opening move during adaptation, in the absence of subsequent mutations, the populations were still highly susceptible to both the drugs. Simultaneous acquisition of specialist mutations to both drugs could have allowed the population ascend the fitness peak along the most direct path. However, this path takes a very long time because it relies on the improbable event of a beneficial double-mutant arising in the evolving population. By contrast, mutational switch-backs provided the population with the opportunity to rapidly and steadily evolve to the combination in an efficient, one-mutation-at-a time, manner (Fig. 5 C).

Having identified the successful evolutionary trajectories leading to resistance, we can re-examine the extent to which the specific selection regimes used may have over or underestimated the efficiency of adaptation by the population in response to the two drugs. Reconstruction of the successful steps to resistance (Fig. 4-5) allows us to speculate on what the most efficient (i.e., fastest) selection regimes could have been in the sequential, as well as the combinatorial studies. For example, Fig. 2 A shows that both the mono-evolved populations made smaller increments during the early days of evolution and made larger increments in tolerable drug concentration towards the end. This implies that the populations were potentially tolerant to higher drug concentrations at an earlier time point and thus the selection gradient could have been increased more steeply. LVS evolving to DOX alone accumulated three mutations relevant to resistance – mutations in *fupA/B, acrAB-tolC* and 30S ribosomal subunit genes. The three mutations together imparted a DOX MIC of 8 µg/ml, i.e. it was tolerant to at least 4 µg/ml of DOX. In Fig 2 A, the DOX mono-evolved populations (dark orange line) were growing at 4 µg/ml DOX around day 40, which is also around the same time the population was able to make large leaps in drug concentration. This implies that the selection environment could have been swapped to CIP at that point. Once the environment was swapped, the population needed one additional mutation in *gyrA* to become CIP resistant. From Fig. 2A, we can see that when the condition was swapped, the population progressed very slowly during the earlier days when low concentrations of CIP were present. Around day 75 ([CIP] = ∼0.2 µg/ml) the population was able to make larger leaps in CIP concentration. It is likely that the gyrase mutation arose in the population at that time and reached fixation by day 75 (Fig. 2A). If we were to adjust the timeline of sequential evolution to eliminate all the days spent by the population in each drug after the relevant mutations had fixed, we estimate that ∼65 days of evolution would be the adaptive timeline (40 for DOX mono-evolution and 25 for sequential evolution to CIP). For sequential evolution of CIP^R^ populations to DOX, this would be ∼45 days (25 for CIP mono-evolution and 20 for sequential evolution to DOX). From the frequency plot shown in Fig. 5B for the population undergoing adaptation to the drug combination, we can see that on day 37 of the experiment, all 4 mutations had fixed in the population. This is substantially shorter than the postulated minimum times taken for fixation of mutations in the sequential populations. Although speculative, this analysis suggests that the drug combination did not delay resistance, and likely accelerated it.

Several important questions remain to be answered. First, if gyrase mutations could confer the pleiotropic effect that FupA/B mutations did, why did gyrase mutations not appear as an opening move in any of the evolving populations? At least some of the FupA/B mutations observed in this study were clearly loss-of-function mutations (Fig. 3 A) while mutations to gyrase are made up of specific SNPs. There are many more ways to inactivate a protein than produce specific changes at the active site conferring enhanced properties and thus, the accessible number of adaptive FupA/B mutations may have produced a strong mutation supply bias in their favor despite their smaller selective advantage, as has been seen in other systems (38, 39). This is supported by the observation that there were more unique adaptive mutations in FupA/B than gyrase (18 vs. 5, shown in Fig. 3 A). Second, it is curious that gyrase mutations never made an appearance in the DOX mono- selected populations, but FupA/B mutations did. Third, why would a gyrase mutation that alters the target of CIP in the cell, confer resistance to DOX, which inhibits an entirely different cellular machinery?

This study provides the genetic basis for the evolution of resistance during exposure to a drug combination and underscores the importance of studying evolutionary dynamics of drug combinations to evaluate their efficacy. It remains to be determined what specific selection conditions may strongly favor mutational switch-backs. As a phenomenon that accelerates evolution of resistance for non-interacting drug pairs and leads to the more rapid acquisition of multi-drug resistance, we should develop a clear understanding of the conditions leading to mutational switch-backs and seek to avoid them in clinical settings. This study was performed at sub-MIC levels throughout and thus one might wonder whether such a condition is found *in vivo* during combinatorial therapy. It is well established that during an infection, pathogens in the host do not experience the same drug concentration across all body compartments and tissues. Spatial and temporal heterogeneity in drug concentration can lead to sub-inhibitory conditions, and subsequent migration of the agent from one host niche to another can also lead to variations in the drug concentration received and mutations acquired (16,40–42). Especially in cases where a combination of drugs is applied, bacteria in different host niches may not only experience sub-inhibitory concentrations but may also experience one drug more than the other. In such cases, mutational switch-backs could facilitate the acquisition of multi-drug resistance. This theory has been modelled in the case of combination drug therapy during HIV infection where resistance to multiple HIV drugs occurred in a sequential and predictable manner (16, 43). A better understanding of how drug combinations affect adaptation to multi-drug resistance could allow clinicians to make more informed decisions on the treatment of patients for which acquisition of antibiotic resistance is a serious concern (44).

## Materials and Methods

### Strains and growth conditions

*Francisella tularensis* subsp. *holarctica* live vaccine strain (LVS) was kindly provided by Dr. Karen Elkins. Strains and plasmids are listed in Table 1. Liquid cultures were routinely grown in BHIC - BHI (BD® 211060) supplemented with 0.1% L-cysteine-hydrochloride (Sigma-Aldrich C1276) and shaken at 37°C, 225 rpm for 2-3 days. BHI powder was dissolved in de-ionized water according to manufacturer’s instructions and sterilized by autoclaving. A stock solution of 10% L-cysteine hydrochloride was made in de-ionized water, filter sterilized and stored at 4°C.

This was added to BHI broth just before use at 0.1% final concentration. Agar plates made of Cystine Heart Agar (CHA, Research Products International C40010-500.0) supplemented with 1% hemoglobin (Thermo Scientific R451402) were prepared following manufacturer’s instructions. When specified, BHI+0.1% cysteine hydrochloride supplemented with 15 g/l agar (BHIC agar) served as solid growth medium. Agar plates were incubated at 37°C for 2-4 days to allow growth. Doxycycline hyclate (TCI Co. Ltd., D4116) was dissolved in de-ionized water and filter sterilized before use. Ciprofloxacin (TCI Co. Ltd., C2227) was dissolved in 0.1N hydrochloric acid and filter sterilized before use. Pre-made stocks of antibiotics were stored at - 20°C for up to 1 week.

For allelic replacement, NEB® 5-alpha *Escherichia coli* competent cells (C2987H) were used for cloning. Transformed *E. coli* cells were selected on LB Miller (IBI Scientific IB49040) supplemented with 15 g/l agar and 50 µg/ml Kanamycin sulfate (Bio Basic KB-0286) at 37°C. Transformed LVS cells were selected on BHIC agar supplemented with 5 µg/ml Kanamycin sulfate at 37°C. For sucrose counterselection, BHIC agar was supplemented with 10% sucrose (50% stock solution made by mixing 125g of sucrose (Bio Basic SB0498) in water to make up 250 ml and sterilized by autoclaving) and counter selection was done at 30°C.

### Minimum inhibitory concentration (MIC) assays

For the checkerboard assay, 2-fold gradients of doxycycline and ciprofloxacin in 100 µl BHIC were created along each axis of a sterile 96 well plate. Wells were inoculated with 5 µl of LVS cultures normalized to OD_600_ 0.05. Outermost wells on the plate were filled with plain media to avoid evaporation of liquid from the assay wells. Plates were grown by shaking at 225 rpm at 37°C for 48 hours in a shaking incubator or in a plate reader shaking orbitally at 282 cpm at 37°C for 48 hours (BioTek Epoch2).

Agar dilution MIC assays were performed in 100 mm diameter petri dishes, as explained in (17), with a few changes. Petri dishes contained BHIC agar and two-fold dilutions of antibiotics. Overnight cultures of LVS isolates (in biological triplicate) growing in BHIC broth were diluted to OD_600_= 0.05 in a 96 well plate. A 96 pin applicator was used to spot the cultures on the agar plates. Spots were allowed to dry before plates were inverted and incubated at 37°C for 2 to 4 days. A no-drug control plate and the ancestor LVS strain were included with each assay.

### Experimental evolution

LVS was evolved using the serial flask transfer approach. 5 biological replicate populations were maintained for each condition. Evolution conditions included mono-selection to doxycycline or ciprofloxacin, sequential selection and combinatorial selection. The control experiment was to serially passage LVS in BHIC broth with no drugs.

For mono-selection, glycerol stock of LVS was streaked onto a CHA+1% hemoglobin plate and grown for 2-3 days. 10 ml cultures in BHIC were started by inoculating each tube with 1 isolated colony from the streaked plate. Tubes were shaken for 2-3 days until confluent growth was visible. 1% or 100 µl of the culture was transferred to a tube containing 10 ml of fresh BHIC. Each passaged population was preserved as a 15% glycerol stock in three cryogenic tubes, 1.5 ml each. For the next passage, populations were transferred to tubes containing a sub-inhibitory concentration of the drug that they were being evolved to. Following this, every passage involved transfer of 1% population to 3 tubes, one containing the same drug concentration as the tube that showed confluent growth and two tubes containing slightly higher drug concentrations, which were empirically determined based on the performance of the populations. Adaptation was stopped when populations evolving to ciprofloxacin grew at at-least 1 µg/ml and populations evolving to doxycycline grew at at-least 8 µg/ml. For combinatorial selection, the experiment was initiated the same way as above with both drugs being introduced simultaneously in a 1:8 ratio (CIP:DOX). Evolution was carried out until the populations grew at at-least 1 µg/ml ciprofloxacin + 8 µg/ml doxycycline. These concentrations were selected because they were at least 2 fold higher than the CLSI breakpoint of these drugs for *Francisella* (29).

For sequential selection, populations evolved to each drug individually were further evolved to the second drug i.e. ciprofloxacin evolved populations were evolved to doxycycline and vice versa. Glycerol stocks of the five replicate populations evolved to drug 1 were thawed and 100 µl of each were mixed and diluted in BHIC broth to 3ml volume. This mixture became the ancestor for sequential evolution of drug 1 resistant populations to drug 2 (If drug 1 is ciprofloxacin, drug 2 is doxycycline and vice versa). Four conditions of sequential evolution were used: (i) Outgrowth of drug 1 resistant population in plain BHIC (ii) Growing drug 1 resistant population at a constant concentration of drug 1 which was the highest concentration to which it was evolved (iii) Evolving drug 1 resistant population to drug 2 (iv) Evolving drug 1 resistant population to drug 2 in the presence of constant concentration of drug 1 (which is equal to the highest concentration at which the starting population grew in the preceding mono-selection experiment). Constant concentration of ciprofloxacin used was 1 µg/ml and doxycycline was 8 µg/ml.

At the end of evolution, each evolved population was serially diluted and spread on CHA + 1% hemoglobin plates to obtain individual colonies termed as end point isolates. 2 to 4 end point isolates were selected from each population. Each isolate was colony purified, grown in BHIC broth (no antibiotics) and frozen as a 15% glycerol stock for long term storage.

### Allelic replacement

Allelic replacement in LVS was carried out using the suicide vector pMP812 (kindly provided by Dr. Martin S. Pavelka) (45) as described in (46), with minor changes. 1000 bp sequences flanking the allelic replacement site on the LVS genome were cloned in pMP812 by Gibson Assembly® (New England BioLabs Inc.) using primers listed in Table 2. LVS cultures were grown in BHIC, single cross over candidates were selected on BHIC agar + 5 µg/ml kanamycin plates at 37°C and sucrose counterselection was done on BHIC agar + 10% sucrose plates at 30°C.

**Table 2:**
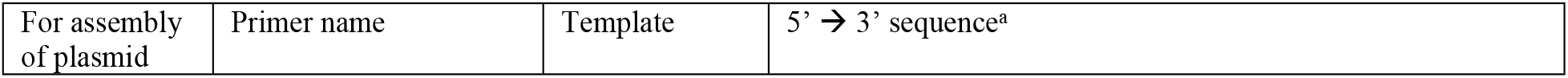
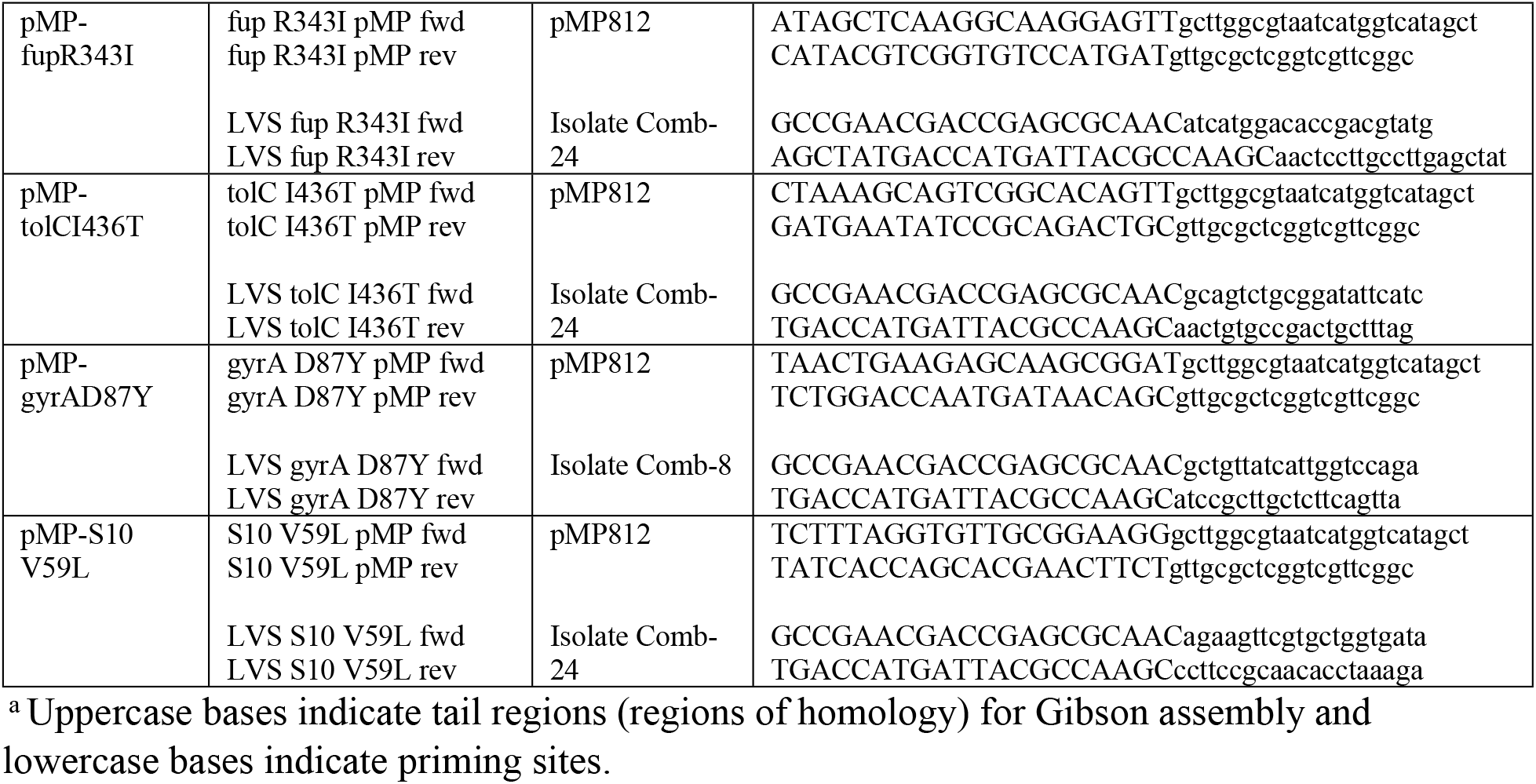
Gibson assembly primers

### Whole genome sequencing and mutation analysis

Final day populations for each evolution condition were sequenced along with 2 to 4 end point isolates from each population. Each condition had 5 replicate populations with the following exceptions - 3 replicate populations for doxycycline resistant ancestor outgrowth in BHIC, 4 replicate populations each for ciprofloxacin resistant ancestor outgrowth in BHIC and ciprofloxacin resistant ancestor evolving to doxycycline. Missing populations were lost to contamination. For populations undergoing combination evolution, each daily population was also sequenced (The populations were sampled only on days when passaging was conducted). The ancestor LVS strain was also re-sequenced.

Genomic DNA from all the samples was extracted using Qiagen DNeasy Ultracean Microbial Kit (Cat No./ID: 12224) following manufacturer’s instructions. Final day populations of samples evolved to doxycycline alone, ciprofloxacin alone and to no drug were sent to a commercial facility for library preparation and whole genome sequencing with at least 200X coverage on the Illumina HiSeq platform to obtain 2 x 150 bp reads. For all other samples, Illumina compatible libraries were prepared using plexWell^TM^ 384 NGS multiplexed library preparation kit from seqWell^TM^. Pooled libraries were sent to a commercial facility for whole genome sequencing with at least 200X coverage on the Illumina HiSeq platform to obtain 2 x 150 bp reads.

All raw fastq reads were trimmed using Sickle (47). Short reads from the re-sequenced LVS strain were aligned to the whole genome sequence of LVS obtained from NCBI (NCBI Reference Sequence: NC_007880.1) using the software breseq version 0.33.2 (48) and identified differences were applied to the reference sequence using the APPLY function in breseq (gdtools) to obtain a modified reference. This modified sequence was used as the reference sequence for all subsequent comparisons. All populations were run using the polymorphism flag -p with the default 5% cut-off frequency. All end point isolates were run using the default consensus mode. The NOTS cluster run by Rice University’s Center for Research Computing (CRC) was used to run these operations (This work was supported in part by the Big-Data Private-Cloud Research Cyberinfrastructure MRI-award funded by NSF under grant CNS-1338099 and by Rice University).

### Tetracycline and ciprofloxacin accumulation assay

Tetracycline was used as a proxy for doxycycline to take advantage of the colorimetric assay that could be performed with tetracycline but not doxycycline. Doxycycline belongs to the tetracycline class of antibiotics and strains used for this assay had the same level of resistance to tetracycline as they did to doxycycline. Tetracycline accumulation assay was carried out as described in (49) with minor modifications. LVS and mutant strains were grown for 48 hours in BHIC and a cell mass equivalent to 5 OD units was used per assay. Cells were exposed to 1000 µg/ml tetracycline for 15 minutes. Ciprofloxacin accumulation was assayed as per Method F described in (50) with minor modifications. LVS and mutant strains were grown for 48 hours in BHIC following which a cell mass equivalent of 20 OD units was used per assay. Cells were exposed to 50 µg/ml ciprofloxacin for 5 minutes. Wet weight of cell pellets was used to normalize accumulated antibiotic concentrations. Assays were conducted in biological triplicate.

## Acknowledgments

The authors would like to thank Dr. Karen Elkins for kindly providing the live vaccine strain (LVS) and Dr. Martin Pavelka for the suicide vector, pMP812. The authors would also like to extend gratitude to Dr. Bruce Levin for critical reading of the manuscript. Funding: This work is supported by funds from the Defense Threat Reduction Agency (grant HDTRA1-15-1-0069) to Y.S. C. Miller is supported by NIH grants P20GM104420 and GM076040. The content of the information in this paper does not necessarily reflect the position or the policy of the federal government, and no official endorsement should be inferred.

## Author contributions

Y.S. conceptualized the idea. C. Marx and C. Miller provided critical feedback. H.H.M. performed the experiments. D.I. assisted in preparing plasmids for allelic exchange. H.H.M., C. Miller, C. Marx and Y.S. analyzed the data and wrote the manuscript.

## Data availability

The whole-genome sequencing data generated during this study were submitted to the Sequence Read Archive (SRA) database under BioProject accession number PRJNA669398.

## Supporting information

S1 Text: Conserved order of appearance of mutations

Figure S1: Consistent parallel evolution observed among replicate populations undergoing combinatorial selection

Table S1: End point isolates obtained after mono-selection, MIC values and mutant alleles identified in genes implicated in resistance

Table S2: End point isolates obtained after sequential selection of doxycycline resistant population, MIC values and mutant alleles identified in genes implicated in resistance

Table S3: End point isolates obtained after sequential selection of ciprofloxacin resistant population, MIC values and mutant alleles identified in genes implicated in resistance

Table S4: End point isolates obtained after combinatorial selection, MIC values and mutant alleles identified in genes implicated in resistance

Data set S1: Mutations identified among evolved populations and end point isolates during mono-selection

Data set S2: Mutations identified among evolved populations and end point isolates during sequential-selection of DOX resistant population to CIP

Data set S3: Mutations identified among evolved populations and end point isolates during sequential-selection of CIP resistant population to DOX

Data set S4: Mutations identified among evolved populations and end point isolates during combinatorial selection to CIP and DOX

Data set S5: Longitudinal mutational data of five replicate populations evolving to combination of CIP and DOX

